# Microbubble Backscattering Intensity Improves the Sensitivity of Three-dimensional (3D) Functional Ultrasound Localization Microscopy (fULM)

**DOI:** 10.1101/2025.04.16.649234

**Authors:** YiRang Shin, Qi You, Yike Wang, Matthew R. Lowerison, Bing-Ze Lin, Pengfei Song

## Abstract

Functional ultrasound localization microscopy (fULM) enables brain-wide mapping of neural activity at micron-scale resolution but suffers from limited sensitivity due to sparse and noisy microbubble (MB) detections. Extending fULM into three dimensions (3D) further exacerbates these challenges because of low-frequency matrix arrays, reduced localization efficiency, and severe data sparsity. To address these limitations, we developed a statistical framework that models MB arrivals in 3D as a Poisson process accounting for localization efficiency, detection probability, and backscattered amplitude. This analysis predicts that integrating amplitude with count-based fULM improves functional sensitivity, particularly under high MB concentrations where localization saturates. Three-dimensional MB advection simulations confirmed these predictions, showing that backscattering fULM (B-fULM) maintains sensitivity at higher MB concentrations where conventional fULM fails. In rat brain experiments, B-fULM yielded stronger and more robust stimulus-evoked responses, with SNR gains of 18% in the somatosensory cortex and 61% in the thalamus, while preserving super-resolved spatial detail (33.4 μm for B-fULM vs 35.7 μm for fULM). These results establish B-fULM as a practical and sensitive approach for super-resolved 3D functional neuroimaging

## I. INTRODUCTION

The brain’s blood supply is precisely regulated by the microcirculation to adapt continuously to local neural activity and metabolic demands [1]. This regulation governs how hemodynamic responses propagate through the vascular network, spanning from deep capillaries to surface pial arteries. While conventional neuroimaging modalities have advanced our understanding of brain structure and functional organization [2–4], mapping the whole-brain hemodynamic response at the microvascular level remains challenging. For instance, functional MRI (fMRI), while capable of deep brain imaging, often lacks the necessary spatial resolution and is sensitive to motion when used with behaving subjects. Similarly, EEG offers high temporal resolution but has limited spatial specificity. Optical techniques achieve high resolution but are constrained by shallow penetration depth or invasive access to target deep regions [5, 6].

In the past two decades, functional ultrasound (fUS) has emerged as a versatile neuroimaging modality, offering a unique combination of high sensitivity, high spatiotemporal resolution (~100 *µ*m spatial, ~1 Hz temporal), and deep tissue penetration [7, 8]. fUS maps brain activity by detecting subtle changes in cerebral blood volume (CBV), with applications spanning animal models [9] to humans [10, 11]. However, its mesoscopic resolution is insufficient to resolve individual microvessel compartments. As shown in Fig. 1c, fUS detects bulk hemodynamic responses and functional changes appears spatially diffuse across larger vascular territories. Yet it is at the microvessel level where neurovascular responses are initiated [12] and where many pathologies originate before propagating to larger vessels [13].

**Figure 1.**
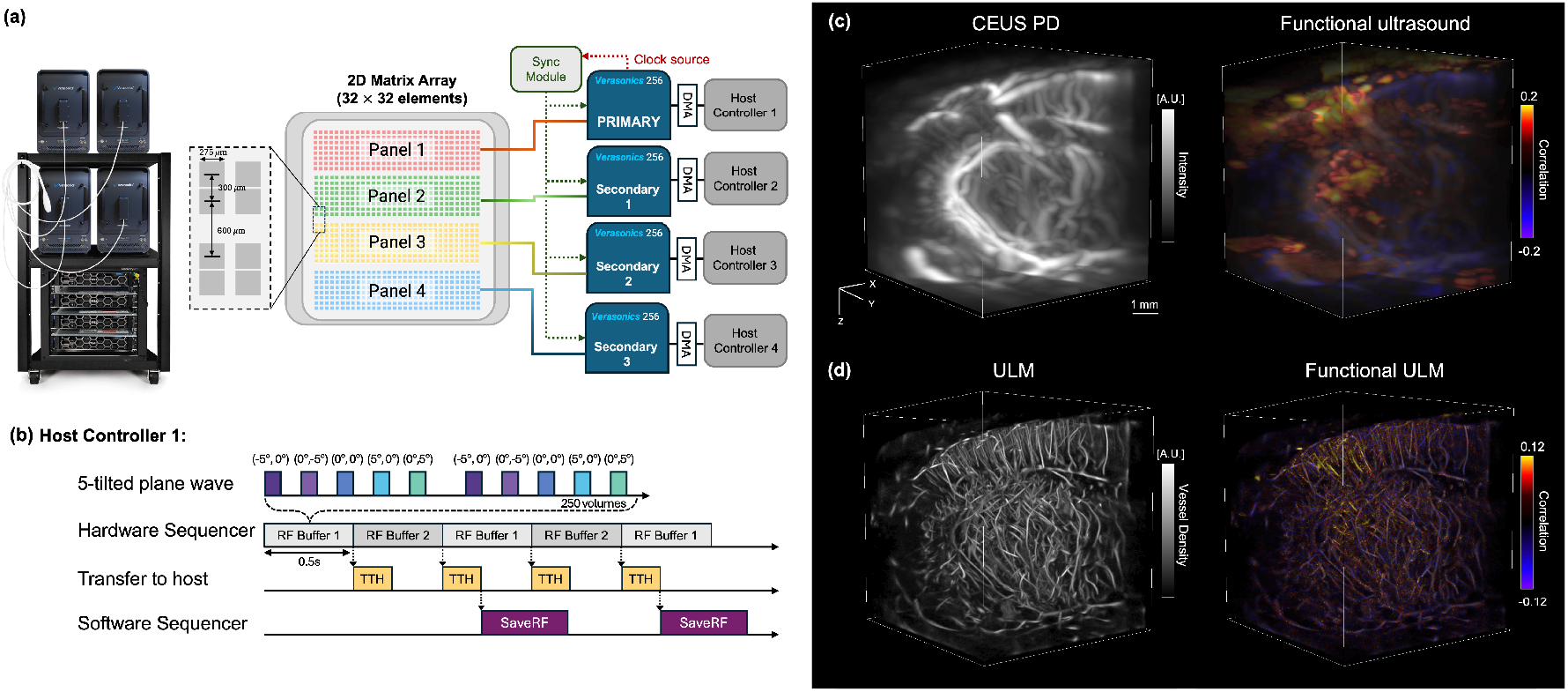
3D functional ultrasound acquisition and imaging pipeline. **(a)** A 32×32 matrix array (8 MHz, Vermon, Tours, France) was driven by a 1024-channel ultrasound system (Verasonics NXT). The array was divided into four subpanels (8×32 elements each), with each panel independently controlled by one NXT system and its host PC. **(b)** Volumetric plane-wave transmissions were performed using five steering angles (−5°, 0°), (0°, −5°), (0°, 0°), (0°, 5°), and (5°, 0°). Each RF buffer contained 250 compounded volumes (~0.5 s acquisition at 500 Hz). A dual-buffer strategy enabled continuous acquisition, with raw RF data transferred to the host PC and saved locally to SSDs. (**c)** Left: 3D contrast-enhanced power Doppler showing vascular anatomy; Right: 3D functional ultrasound activation map via voxel-wise correlation with stimulus pattern. **(d)** Left: 3D ULM vessel density reconstruction; Right: 3D functional ULM activation map correlating MB flux with the stimulation pattern.

Ultrasound localization microscopy (ULM) is a super-resolution technique that surpasses the acoustic diffraction limit. By localizing and tracking individual microbubbles (MBs) in circulation, ULM generates vascular maps with micron-scale resolution [14]. A key advantage of ULM is its ability to provide quantitative metrics of blood flow, including velocity [15] and direction [16]. The primary drawback, however, is its low temporal resolution. Fully reconstructing small vessels (~5 − 10*µ*m) requires minutes of data accumulation [17], limiting its application to static vascular imaging.

Recently, functional ULM (fULM) was introduced to detect task-evoked hyperemia in the rodent brain, achieving ~6.5*µ*m spatial resolution with ~1s temporal resolution [14]. fULM uses MB flux (the count of localized MBs per pixel per time) as its functional readout. As illustrated in Fig. 1d, fULM enhances spatial resolution by resolving individual microvessels and their localized task-evoked responses. Compared to conventional fUS, fULM provides distinct advantages. It can disentangle compartment-level vascular responses [18], quantify absolute vessel parameters such as diameter and flow velocity at the level of individual vessels [19], and detect early microvascular dysfunctions that precede macroscopic changes [20]. These capabilities offer insights into neurovascular coupling and disease mechanisms that remain inaccessible with bulk CBV measurements from fUS or fMRI. However, because the MB detection events are sparse at small vessels, repeated stimulus trials (often over 10 trials) are needed to resolve capillary-level flow changes [14]. As a result, fULM inherently trades off spatial resolution and sensitivity against temporal resolution.

One natural strategy to reduce repetitions is increasing the MB concentration, thereby increasing the number of MB detections per trial. However, elevated MB concentrations exacerbate MB PSF overlap, degrading the localization accuracy. As MBs merge spatially, localized counts saturate and underestimates true MB flux −even when actual MB flux continues to increase [21]. Recent deep learning-based ULM methods have shown that improved localization under high MB concentrations can restore vessel filling and enhance fULM activation with fewer repetitions [22].

Extending fULM into three dimensions (3D) poses an even greater challenge. Current 3D ULM implementations for small animal imaging often use a 2D matrix array transducer (e.g., the 8MHz, 32×32 matrix array made by Vermon S.A., Tours, France) with ultrafast planewave imaging [23, 24]. The available 2D matrix array transducers operate at lower frequency than their 1D counterparts (e.g., 8 MHz vs. 15 MHz). The lower frequency increases the MB PSF size, making spatial overlap more likely. In addition, their smaller aperture and larger pitch further degrade image quality, introducing spatial aliasing and reducing localization accuracy in 3D. Volumetric super-resolution further exacerbates data sparsity: Interpolating by a factor of *K* in each of three dimensions reduces the average MB arrival rate per voxel by 1/*K*^3^ (compared with 1/*K*^2^ in 2D). Consequently, dividing the volume into 3D voxels dilutes the MB occupancy, making the functional signals even sparser and limiting the sensitivity of 3D fULM. In addition, existing 3D ULM implementations typically acquire 2−3 GB of data per second, which translates to ~2TB of raw RF data per single 3D fULM experiment (15 repetition cycles). Processing such volumes on a typical workstation can take on the order of several days, and these technical barriers have so far prevented demonstration of 3D fULM to date.

Moreover, in fUS, red blood cell (RBC) backscattering intensity has proven to be a reliable indicator of blood volume variations [25–27]. The functional sensitivity of fUS is primarily determined by the signal-to-noise ratio (SNR) of RBC backscattering, which can be enhanced through extensive frame averaging with ultrafast plane wave imaging. However, conventional fULM use MB count as an indicator of blood volume variations which is subject to numerous confounding factors, including MB concentration, localization accuracy, and accumulation time. In cases of increased MBs due to functional hyperemia, the backscattering intensity of MBs may serve as a more effective indicator of blood flow changes than MB count. This effectiveness of MB backscattering intensity in fUS was demonstrated by Errico *et al*. in 2016 [28]. Additionally, Renaudin [29] highlighted that leveraging the backscattering amplitude of MBs enhances the sensitivity of 2D backscattering structural imaging compared to conventional 2D ULM imaging by recovering information missing in the out-of-plane direction. Motivated by these findings, we introduce a novel method that combines MB back-scattered amplitude with localized MB positions to improve functional sensitivity of 3D fULM without compromising spatial resolution.

The remainder of this paper is organized as follows. Section II develops the statistical framework for 3D fULM and B-fULM, deriving closed-form expressions for functional sensitivity and introducing 3D MB advection simulations that validate the theoretical predictions. Section III presents results from the simulations and *in vivo* rat brain experiments under whisker stimulation. Finally, Sections IV and V provide a discussion of findings and conclusions of this study.

## II. METHOD

### A. Modeling MB Signal Statistics

Let the beamformed image be interpolated by a factor of *K* along each of the *n* dimension (each voxel is split into *K*^*n*^ equal super-resolved voxels). Let *γ*_0_ be the mean number of true MB arrivals per frame in an original (pre-interpolated) voxel. Then the mean arrival rate per super-resolved voxel is give as,

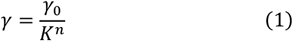

Within each super-resolved voxel, true MB arrivals per frame follow a Poisson process:

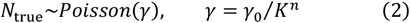

After interpolating by a factor of *K* in each of *n* dimensions, arrivals in each super-resolved voxel remains Poisson with mean *γ* = *γ*_0_/*K*^*n*^. Thus in 3D (*n* = 3) the per-voxel mean is diluted by 1/*K*^3^ (vs. 1/*K*^2^ in 2D), lowering voxel occupancy and increasing MB signal sparsity. Because of the limited spatial resolution and MB overlap, only a fraction of MB arrivals is isolated and localized as distinct positions. Let *N*_*loc*_ be the number of localized MBs and *π*(·) ∈ [0,1] be the localization fraction:

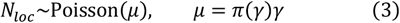

where 0 ≤ *π*(*γ*) ≤ 1 is the localization probability (fraction). At low-concentration, *π*(*γ*) is close to 1, but at higher concentrations *π*(*γ*) decreases due to MB overlap. In the derivations below, we consider small functional contrasts and take *π*(*γ*) as approximately constant and represent localization fraction at baseline flow as *π*_0_ = *π*(*γ*_0_).

To account for the MB signal that is captured even when bubbles merge (i.e., contribute backscattered-intensity to some localized position), let each true MB be captured with detection probability *ρ*(*γ*) ∈ [0,1]. Then

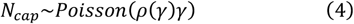

Assuming each captured MB contributes a constant amplitude of *a*, the localized intensity per frame is

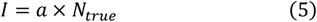

The expected intensity and variance follow the Poisson model of *N*_*true*_:

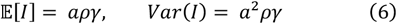

Importantly, this intensity-based measure depends on the captured MBs *N*_*cap*_ (via *ρ*(*γ*)), whereas the count *N*_*loc*_ is further reduced by the subset that is localized. Typically, under MB crowding, *ρ* ≥ *π*, as overlapping MBs form fewer but brighter detections.

### B. Functional Sensitivity Derivations

Using the statistical model above, functional sensitivity can be quantified by a z-score under averaging *M* independent frames. Let the baseline MB arrival rate be *γ*_0_, and the activated MB arrival rate be

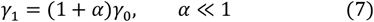

where *α* denotes the fractional increase during activation. We assume that localization and detection probability remains approximately constant over this range, i.e. *π*(*γ*_0_) *≈ π*(*γ*_1_) = *π*_0_, *ρ*(*γ*_0_) *≈ ρ*(*γ*_1_) = *ρ*_0_.

For count-based fULM, defining *µ*_*i*_ = *π*_0_*γ*_*i*_, the z-score is:

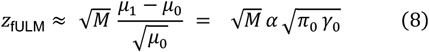

For B-fULM, with 𝔼[*I*_*i*_] = *aρ*_0_*γ*_*i*_ and Var[*I*_0_] = *a*^2^*ρ*_0_*γ*_0_,

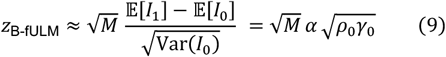

Taking the ratio yields,

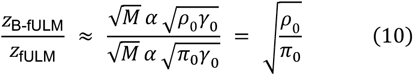

This result shows that B-fULM achieves higher sensitivity than count-based fULM whenever *π*_*o*_ < *ρ*_0_. In the case where all MBs are isolated (*π*_*o*_ = 1, *ρ*_0_ = 1, very dilute), the two methods achieve equal functional sensitivity. In practice, however, *π*_*o*_ *≈* 1 is attainable only at very low MB concentrations (typical ULM operates at *γ*_0_ = 0.005 − 0.01 MB/pixel/frame [17, 30]). Under these conditions, a single stimulation period yields too few MB detections for robust separation of baseline and activation. As a result, robust sensitivity requires increasing *M*, which means longer stimuli or repeated stimulation cycles.

An alternative is to increase MB concentration, thereby increasing *γ*_0_. At low acoustic drive, the detection efficiency *ρ*_0_ remains near unity across a broad range of MB concentrations. Experimental studies confirm that backscattered intensity increases linearly with MB concentration starting from the dilute regime, only saturating at very high concentration (~2‰) [31]. Similarly, attenuation-compensated backscattered intensity measurements with Definity show linear scaling over a wide range of concentrations, with deviations emerging at higher pressures and concentrations due to attenuation and nonlinear oscillations [32]. In the linear regime where detection efficiency remains high (*ρ*_0_ *≈* 1), the relative sensitivity simplifies to

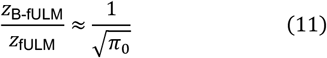

This means that when all arriving MBs contribute to the backscattered signal and completely captured, B-fULM achieves a sensitivity advantage of 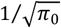 over fULM. In practice, because *π*_0_ decreases rapidly with increasing MB concentration, fULM sensitivity starts to saturate, whereas B-fULM continues to improve linearly until limited by attenuation, saturation, or nonlinear effects.

### C. Simulation for 3D MB Advection and Functional Mapping

We simulated MB advection through a cylindrical vessel embedded within a 3D volume to evaluate functional mapping. Acoustic and sampling parameters were chosen to approximate a 7.8 MHz center frequency (*c* = 1540 m/*s*) with isotropic voxel spacing of *λ*/2 in all three dimensions, sampled at a pulse repetition frequency of 500 Hz. The imaging volume was 49 × 54 × 50 voxels, with the vessel centered and oriented along the lateral axis. MB flow followed a laminar parabolic profile with a partial-slip boundary condition, in which the wall velocity was set to 50% of the centerline velocity. The baseline centerline velocity was 0.05 m/s and was modulated during stimulation.

The stimulation pattern comprised of 31 consecutive blocks of 500 volumes each (1s per block at 500 Hz). The first block served as a baseline, followed by 30 alternating ON/OFF blocks (15 ON-OFF cycles), yielding a total of 15,500 volumes. For each volume, all MBs in a preallocated pool were advanced by one time step; per frame displacement was scaled by *s*_*v*_. New MBs were injected at the vessel inlet according to a binomial spawning process

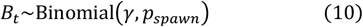

with

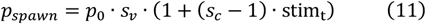

where *p*_0_ = 0.05 is the baseline spawn probability, *γ* is the number of spawning trials, *s*_*v*_ is the velocity scaling, *s*_*c*_ is the concentration scaling, and stim ∈ {0,1} indicates the block state. Newly spawned MBs were assigned inlet positions and velocities were sampled from the laminar velocity profile. MB exiting the volume became inactive and returned to the pool.

To investigate the effect of MB concentration, *γ* was varied from 1 to 15 in unit steps. For each *γ*, five independent realizations were performed to report the mean and standard deviation. The expected MB influx per frame increase linearly with *γ*. At the lowest MB concentration (*γ* = 1), the expected number of newly spawned MB is ~0.05/frame during OFF and ~0.09/ frame during ON state (*p*_0_ = 0.05, *S*_*v*_ = 1.2, *S*_*c*_ = 1.5), corresponding to ~25 and ~45 MBs per 500-frame block. At the highest MB concentration (*γ* = 15), the expected influx increases to ~0.75/frame (OFF) and ~1.35/frame (ON), corresponding to ~375 and ~675 MBs over a 500-frame block.

### D. Experiment setup of the in vivo rat brain imaging study

The *in vivo* rat brain study was approved by the Institutional Animal Care and Use Committee (IACUC) at the Duke University (A012_25_01). A Sprague-Dawley 17-week-old rat was used in this study. A cranial window was opened on the skull of the rat using a rotary Dremel tool. The surgery was implemented under anesthesia using 1.5% isoflurane mixed with medical oxygen. After the surgery, anesthesia was maintained by bolus injection of Urathane (1g/kg). A catheter was inserted into the tail vein of the rat for infusion of the ultrasound contrast agent (DEFINITY®, Lantheus Medical Imaging, Inc.). 0.8 mL of Definity was diluted in 3 mL saline and continuous infusion of contrast was performed at a rate of 30 *µ*L/min using a syringe pump (NE-300, New Era PumpSystems Inc., Farmingdale, NY). Magnetic stir bars with a size of 1mm × 3mm were placed in the syringe and gently stirred during the experiment to stabilize the concentration of the contrast agent. The 2D matrix transducer was mounted on a 3D scanning gantry and positioned on the cranial window of the rat skull. The B-mode and power Doppler images were used as the reference for adjusting the position of the 2D matrix transducer.

Whisker stimulation was used for triggering neural activities in the somatosensory barrel cortex of the rat. As shown in Fig. 2(a), the stimulation protocol included 15 cycles where each cycle included 30 seconds of stimulation (“On”) and 30 seconds of resting (“Off”). In addition, one 30-second resting periods were added before and after the 15 cycles to measure pre-stimulation and post-stimulation brain activities.

**Figure 2.**
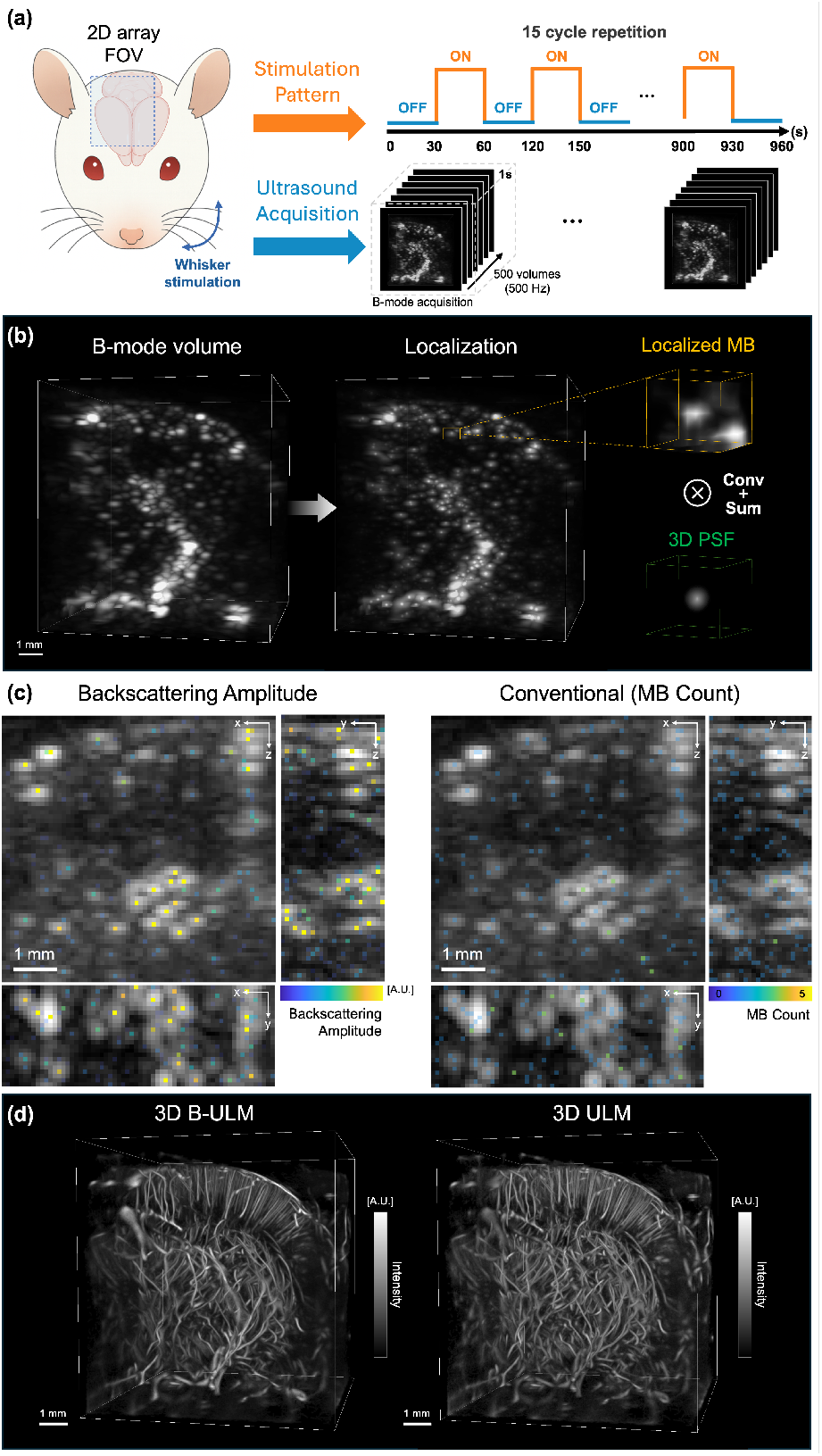
Whole-brain volumetric functional ULM in an anesthetized rat during whisker-evoked activation. **(a)** A 2D matrix array acquires 3D volumes at 500 Hz during 30-s alternating ON/OFF whisker stimulation (15 cycles). (b) After localization, the backscattered signal was estimated by applying a Gaussian-weighted sum of voxel intensities centered at each localized position, and the resulting amplitude was assigned to the corresponding MB detection. (c) Example 2D B-mode projections with MB localizations overlaid, color-coded by backscattering amplitude (left) or by MB count (right). (d) Left: 3D backscattered-intensity ULM vessel density map (B-ULM); Right: 3D ULM density map.

### E. Ultrasound imaging sequences

The ultrasound signal acquisition was synchronized with the stimulation pattern and was acquired using Verasonics NXT multi-system platform (Verasonics, Inc., Kirkland, WA, USA) configured with four synchronized NXT 256 systems (1024 channels). The systems were directly connected to a 32×32 element matrix array (8MHz, Vermon, Tours, France), which is divided into four subpanels. Each subpanel comprised 256 elements arranged in an 8 × 32 grid and was independently controlled by separate NXT systems (Fig 1a). Individual element size is 0.3 mm with a 0.3-mm pitch with 0.6 mm distance between each panel, resulting in a total aperture covering a field of view (FOV) of 9.3 ×10.2 mm.

The matrix probe was operated at a center frequency of 7.81MHz with a pulse repetition frequency (PRF) of 12.5 kHz. Volumetric plane-wave compounding was performed using five steering angles in azimuth/elevation: (−5°, 0°), (0°, −5°), (0°, 0°), (0°, 5°), (5°, 0°), resulting in a post-compounded volume rate of 500 Hz (Fig. 1b). Each RF buffer contained 250 compounded volumes, corresponding to ~0.5s of acquisition at 500 Hz. Raw RF data from each NXT system were transferred to the corresponding host PC and saved locally to separate SSDs. To ensure continuous acquisition, we used a dual-buffer strategy, where a new buffer was initiated immediately as the previous one finished (Fig. 1b). A total of 500 post-compounded volumes were acquired and saved to the host computers within every 1 seconds.

Acoustic output was characterized using a hydrophone (HGL-0085, ONDA) with signals recorded on a digital oscilloscope (TB2000B, Tektronix). The spatial peak of the transmitted beam was identified, and peak negative pressure (PNP) was measured at this location. To match the in vivo experiments, the transmit voltage was set to 8V. At 10 mm depth, the PNP and MI (0.3 dB/MHz derating) were below 57 0.1 MPa and 0.02, respectively.

### F. Ultrasound Localization Microscopy Processing

Following the experiment, raw RF data from all host PCs were combined before beamforming. The beamforming was performed offline using RTX A6000 GPUs (NVIDIA, CA, USA) using a customized CUDA beamforming program. The beamforming resolution was half wavelength per voxel (98.6 *µ*m) in lateral, axial and elevation directions. Tissue clutter was suppressed using singular-value decomposition (SVD) with a fixed low-rank cutoff (rank = 90). To separate flow towards/away from the transducer, the SVD-filtered data were further decomposed using directional filtering [33].

To approximate high MB concentrations, we synthesized combined IQ volumes by summing consecutive acquisition cycles. The experiment consisted of 15 cycles, each containing 60 acquisitions (30 stimulus and 30 rest). For each composite index *k* (*k* = 1, …, 13), a three-cycle window was applied with unit stride,

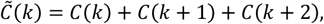

where *C*(*k*) denotes the complex IQ data from cycle *k*. The summation was performed in a time-aligned manner, ensuring that stimulus and rest acquisitions within each 60s period remained synchronized across combined volumes. Consequently, windows (i.e., 1-3, 2-4, …, 13-15) generated 13 composite cycles. These composites were then used to identify the initial MB center by applying voxel-wise normalized cross-correlation (NCC) with a predefined 3D Gaussian kernel approximating the PSF. This was followed by thresholding (≥0.6) and detection of 3D regional maxima as candidate centers [34]. These centers were refined to sub-voxel precision using 3D Farid-Simoncelli interpolation and axis-wise quadratic root finding of first- and second-order derivatives. After localization, MB trajectories were estimated between consecutive volumes using 3D nearest neighbor association, and final 3D ULM maps were reconstructed on isotropic resolution of *λ*/5 (39.4 *µ*m).

Backscattering ULM was generated by first extracting a 3D region corresponding the PSF size around each localized MB. Within this region, the backscattered signal was estimated using a Gaussian-weighted sum of voxel intensities (Fig. 2b), and the resulting amplitude was assigned to the MB detection (Fig. 2c). Backscattering amplitudes were then used to generate amplitude-weighted trajectories that were accumulated into B-ULM maps. B-ULM was reconstructed on the same grid as the ULM maps (*λ*/5, 39.4 *µ*m) (Fig. 2d). 3D functional activation map was computed by correlating the stimulation pattern with either ULM or B-ULM signals, generating fULM and B-fULM maps. The final correlation maps were overlaid with the ULM maps using 3D Slicer.

## III. RESULTS

### A. Simulation Study

Functional sensitivity was evaluated by computing the Pearson correlation between the stimulus pattern and both MB count and backscattering amplitude signals within a fixed intravascular ROI (Fig. 3). In Fig. 3b, spatial activation maps shows that B-fULM correlation increased progressively with MB concentration, maintaining strong and coherent response within the vessel lumen. In contrast, fULM maps showed weak responses across concentrations, without a clear increase at higher MB concentrations.

**Figure 3.**
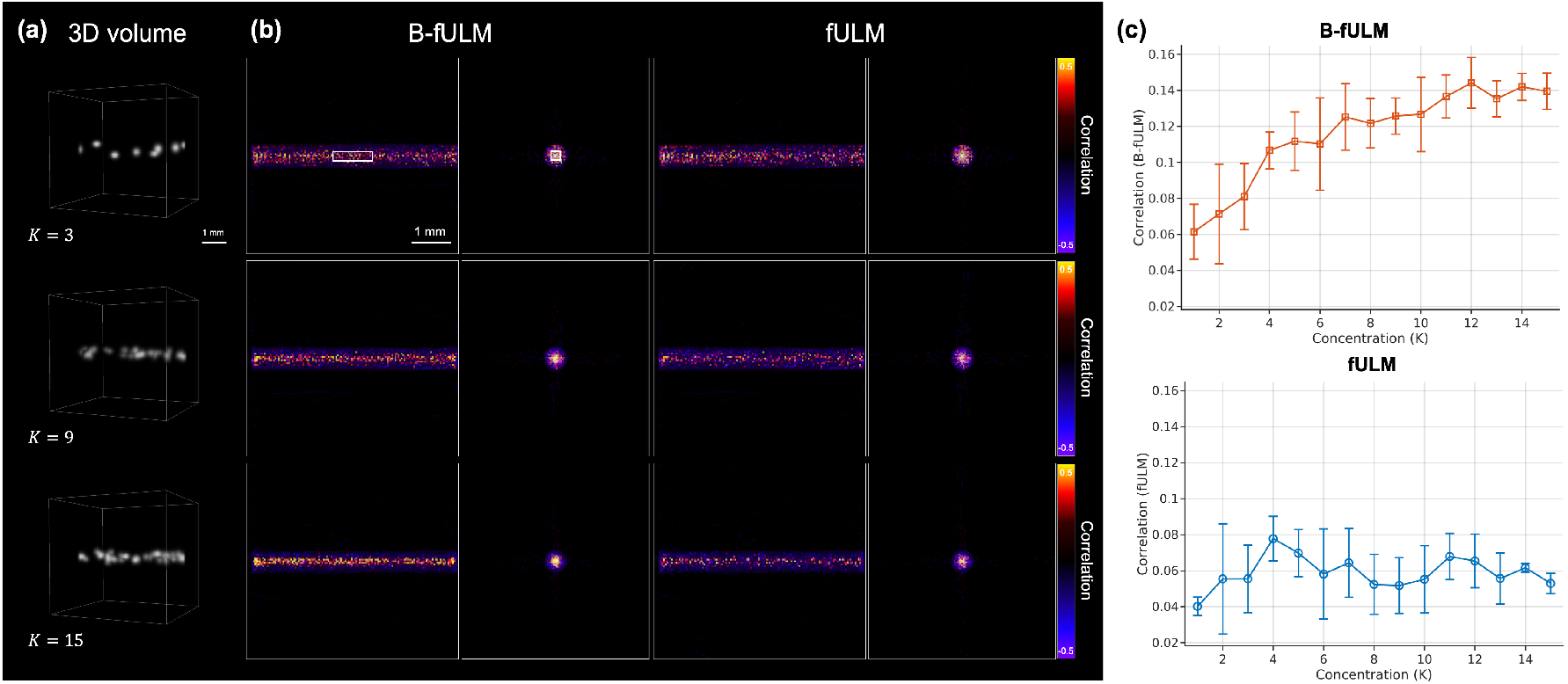
Simulation for 3D MB advection and functional mapping. (a) Representative 3D volumes at increasing MB concentrations (*γ*),showing denser MB population at higher *γ*. (b) Correlation maps for B-fULM (left) and fULM (right) at matched *γ*; with the vessel ROI outlined (white box) (c) Correlation–MB concentration curves (mean ± SD, five repeats per *γ*) for both fULM and B-fULM.

In Fig. 3c, correlation values for B-fULM increased consistently with MB concentration, rising from 0.062 ± 0.015 at *γ* = 1 to a maximum of 0.144 ± 0.014 at *γ* = 12, and shows mild saturation at the upper end (0.135–0.144 range for *γ* = 11 − 15). Across the full range, B-fULM consistently exceeded count-based fULM in correlation values and exhibited a strong increasing trend (slope *≈* 5.35 × 10^−3^, *R* = 0.92), reflecting increasing functional sensitivity at higher concentrations.

In contrast, count-based mapping (fULM) displayed limited a response range (Fig. 3c). Correlation values increased only modestly from 0.040 ± 0.005 at *γ* = 1 to an early maximum of 0.078 ± 0.012 at *γ* = 4. then fluctuated without sustained gain (0.052–0.070 for *γ* = 5 − 15). The fitted slope exhibits a negligible trend (slope *≈* 1.91 × 10^−4^, *R* = 0.094), with a concave overall trend (mean second difference = −1.84 × 10^−3^), indicating poor scalability with increasing MB concentration.

Collectively, these results demonstrate that B-fULM provides stronger activation and functional sensitivity than count-based fULM, with substantial gains at high MB concentration. The enhanced sensitivity of B-fULM implies that reliable detection can be achieved with fewer stimulus repetitions, providing a practical advantage for experimental efficiency. Importantly, these findings are consistent with theoretical predictions: because amplitude integrates the backscattered signal from all MBs, whereas counts rely only on the subset of isolated detections (*π*_0_ < 1), B-fULM retains sensitivity even when MB overlap increases at higher concentrations.

### B. In vivo rat brain study

Figure 4 compares the 3D structural and functional maps obtained with conventional fULM and B-fULM. Both methods recovered highly similar vascular architecture, including major penetrating vessels and deep thalamic branches (Fig. 4a, c). B-fULM exhibited reduced background noise, likely due to lower relative intensity of localized noise signals compared to MB signals. Quantitative Fourier shell correlation (FSC) further confirmed that spatial resolution is preserved (Fig. 5), with B-fULM achieving a resolution of 33.43 *µ*m, slightly finer than the 35.74 *µ*m measured for conventional fULM using a 1/2-bit threshold criterion.

**Figure 4.**
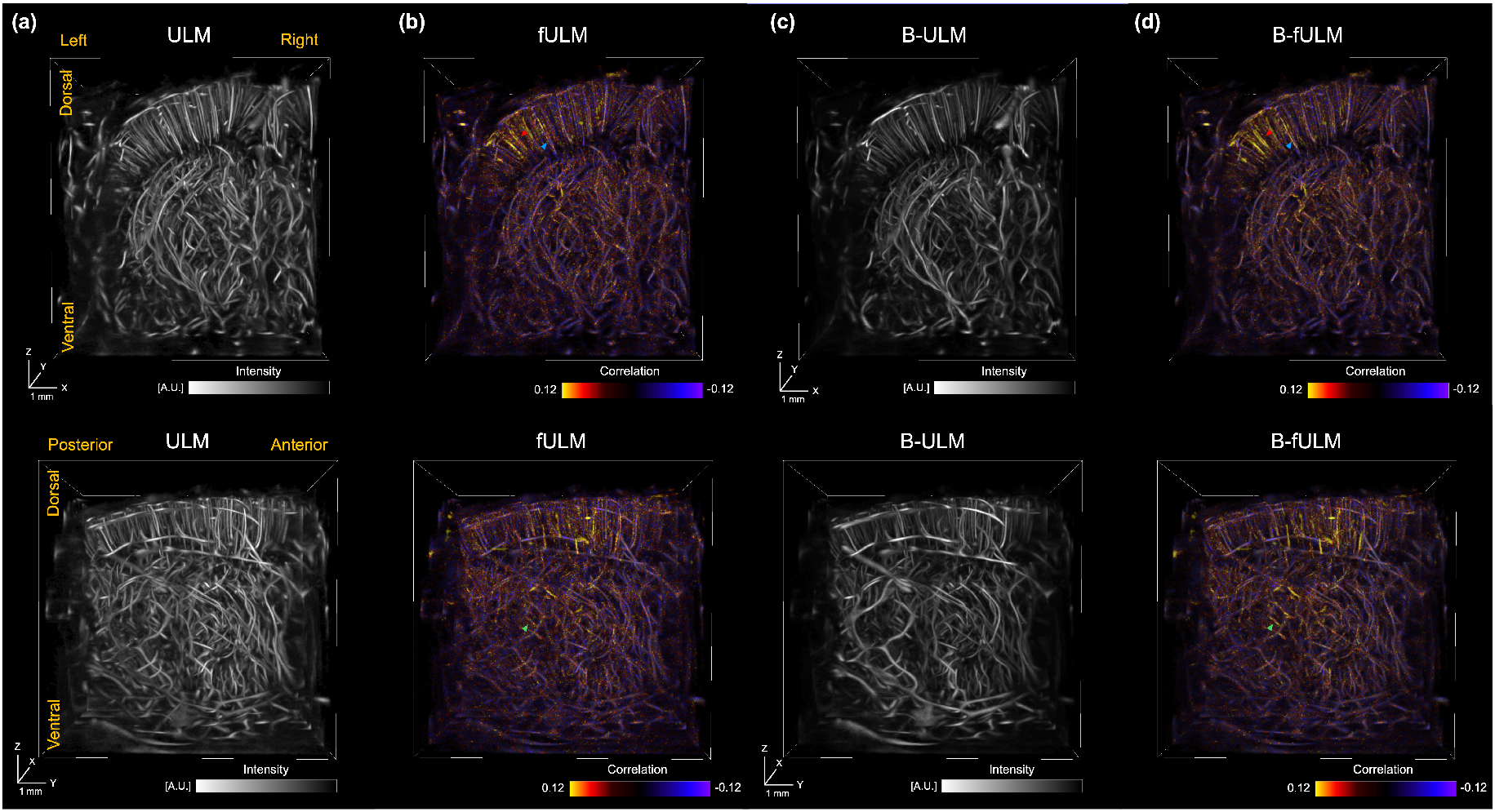
Three-dimensional structural and functional ULM maps of the rat brain under whisker stimulation. **(a)** Conventional ULM structural reconstruction. **(b)** Conventional fULM functional activation map. **(c)** Backscattering ULM (B-ULM) structural reconstruction. **(d)** Backscattering fULM (B-fULM) functional activation map. Top row: dorsal–ventral view. Bottom row: posterior–anterior view.

**Figure 5.**
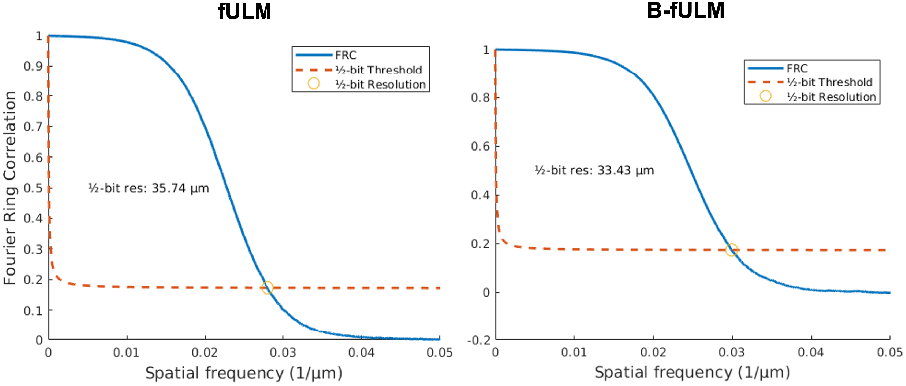
Fourier shell correlation (FSC) analysis of spatial resolution for fULM (left) and B-fULM (right). The solid blue curves represent the measured FSC, while the dashed red lines indicate the 1/2-bit threshold.

In Fig. 4, the functional activation maps produced by B-fULM are notably stronger and exhibits reduced sparsity than conventional fULM. Under whisker stimulation, both methods identified MB flux increase in the somatosensory barrel field (S1BF) cortex and in the ventral posteromedial (VPM) thalamic region, corresponding to the expected neurovascular response. However, the fULM activation map appears sparse and discontinuous where only scattered segments of vessel show correlation to the stimulus (red arrow in Fig 4.b) In the B-fULM map, the activation is stronger and entire vascular segments is highlighted (red arrow in Fig 4.d). In addition, the active vessel traces out the underlying vascular anatomy much more continuously (blue arrow in Fig 4.b, d). B-fULM also shows enhanced signal, whereas fULM shows little or no detected activity (green arrow in Fig 4.b, d). This indicates an enhanced functional sensitivity with B-fULM, yielding dense activation patterns that mirror the underlying structural vasculature.

Fig. 6 illustrates activation results in the S1BF and VPM regions, with zoomed-in comparison of fULM and B-fULM. In Fig. 6 c, d, B-fULM produces more saturated activation maps with broader, more contiguous activation along vessel segments, whereas fULM shows sparse and more fragmented activations. Representative ROIs in S1BF and VPM (white ROIs in Fig. 6c, d) were analyzed to quantify task-induced responses. The cycle-averaged time courses (Fig. 6e, f) show that B-fULM exhibits steeper on/off transitions during stimulus blocks, reflecting strong task-induced responses. In S1BF (Fig. 6e), B-fULM produced stronger functional sensitivity than fULM (|ΔP|: 0.067 vs 0.060; paired t=2.27, p=0.04), corresponding to a ~ 12% larger effect. While B-fULM traces exhibited slightly greater temporal variability (*σ*: 0.047 vs 0.045), the net sensitivity was improved, with SNR enhanced by 18.1% (1.83 vs 1.55). In VPM (Fig 6.f), B-fULM again outperformed fULM, with ~56% increased functional sensitivity (|ΔP|: 0.064 vs 0.041; paired t=3.89, p=0.002) and 61% increased SNR (2.30 vs 1.43). Together, these findings demonstrate that B-fULM achieves higher functional sensitivity than fULM in both cortical and thalamic regions.

**Figure 6.**
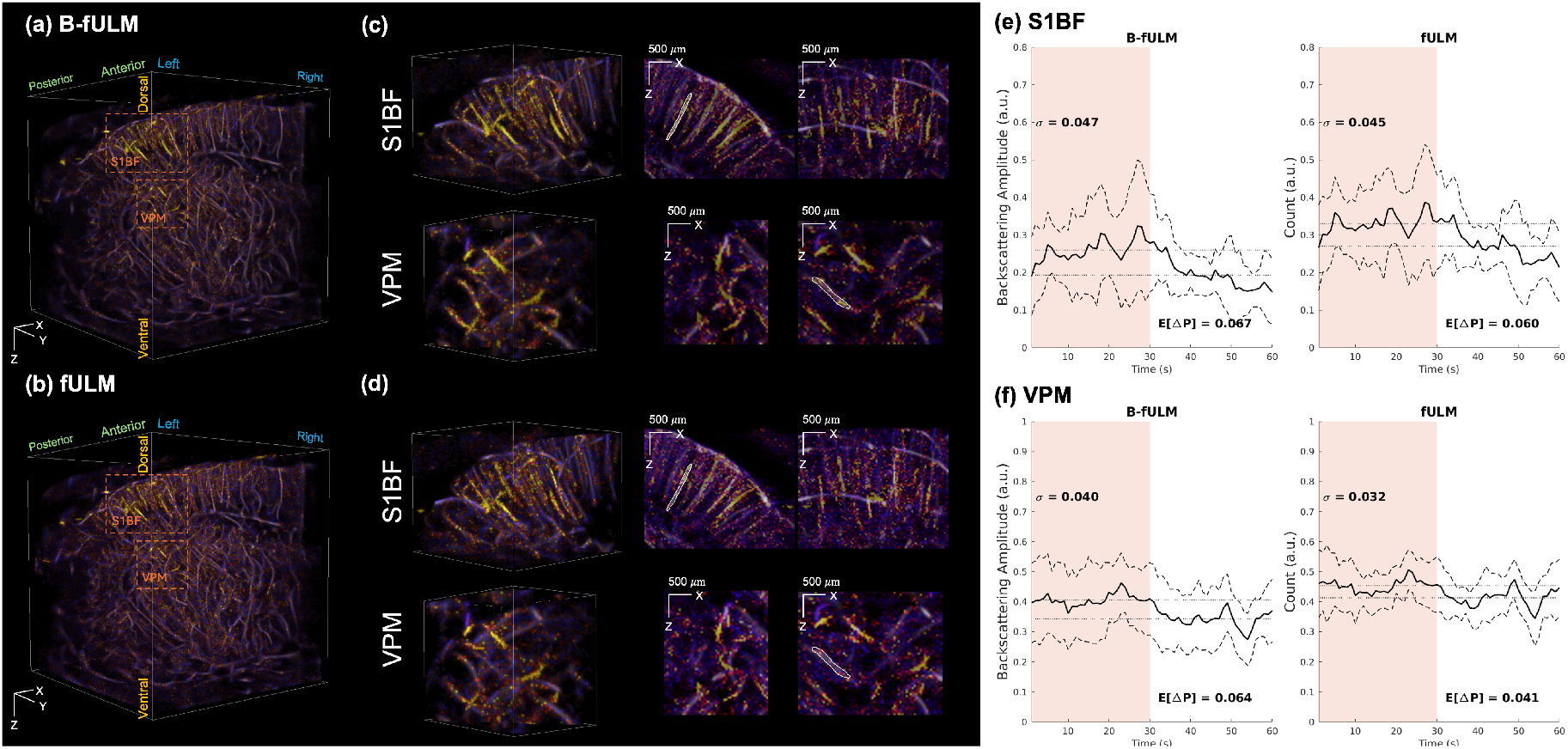
3D functional ULM activation in S1BF and VPM: comparison of B-fULM and fULM. **(a)** B-fULM activation map. **(b)** fULM activation map. **(c**,**d)** Zoomed 3D views of activated vasculature in S1BF (top) and VPM (bottom); panels show maximum intensity projections (MIPs). **(e**,**f)** Cycle averaged functional traces for S1BF (e) and VPM (f) within white ROIs in c,d: B-fULM shows microbubble backscattering amplitude flux and fULM shows microbubble count flux, each averaged across 13-time aligned stimulation cycles. Curves plot the mean (solid) and ± SD across 60s stimulation pattern (dashed); the shaded region denotes the ON period.

## IV. DISCUSSION

While 2D fULM has demonstrated the feasibility of mapping hyperemia with micron-scale resolution, it is limited by narrow spatial coverage and out-of-plane MB motion. Functional responses often span distributed cortical and subcortical networks and sequentially scanning 2D planes cannot fully capture activity across distant regions. In addition, 2D ULM suffers from inaccuracies due to elevation projection, which complicates vessel tracking and flow quantification [35]. In contrast, 3D fULM achieves whole-brain coverage at micron-scale resolution—unattainable with fMRI, optical microscopy, or 3D fUS—allowing vessel-resolved functional mapping across the brain. This enables studies of microvascular contributions to global network dynamics and bridges mesoscopic whole-brain imaging with microscopic optical methods.

Despite these advantages, 3D fULM presents formidable challenges. Compared with 2D imaging, voxel MB occupancy is diluted by cubic interpolation, making functional signals inherently sparse. Lower-frequency 2D matrix arrays further enlarge the PSF, increasing spatial overlap of MBs and reducing localization efficiency. As a result, 3D fULM requires long acquisition times and repeated stimulus cycles to achieve reliable sensitivity, producing massive data volumes on the order of terabytes per experiment. Together, these physical and technical constraints explain why most fULM studies have remained 2D, despite the recognized need for brain-wide 3D imaging. In this work, we addressed these limitations with two complementary innovations. First, we introduced backscattering fULM (B-fULM), which integrates MB backscattering intensity with MB count. By incorporating the full backscattered signal, B-fULM preserves sensitivity gains even when MB concentration increases, overcoming the saturation effect that limits conventional fULM. Theoretical analysis established that B-fULM achieves higher functional sensitivity than count-based fULM except under ideal sparse conditions. 3D MB advection simulation confirmed this prediction, demonstrating that B-fULM correlation increase monotonically with MB concentration, reaching a maximum of 0.144 ± 0.014 at *γ* = 12, while fULM correlations plateau early around ~0.07 (Fig. 3c).

Second, we implemented a synchronized multi-system acquisition platform capable of continuous 1024-channel imaging at high volume rates (500Hz). This configuration allows uninterrupted data acquisition, essential for functional imaging. From the sensitivity equation (Eq 8, 9), increasing the number of volume (*M*) directly enhances detection sensitivity, making continuous acquisition at high volume rates critical. High volume rates further improve vessel reconstruction and velocity estimation by preserving the temporal fidelity of MB trajectories [36]. Continuous acquisition also eliminates dead time between buffers, ensuring that the full stimulus-evoked hemodynamic response is captured and that high-quality 3D mapping of microvascular dynamics is maintained throughout the entire experiment.

*In vivo* whisker stimulation study on the rat brain confirmed the combined advantages of B-fULM and multi-system acquisition. Structural resolution was maintained, with Fourier Shell Correlation (FSC) showing 33.43 *µ*m for B-fULM compared to 35.74 *µ*m for fULM (Fig. 5). In terms of functional sensitivity, B-fULM produced increased responses (Fig. 6): in the S1BF cortex, functional sensitivity increased by 12% (|ΔP|: 0.067 vs. 0.060) with by 18.1% improvement in SNR. In the VPM thalamus, the gain was even larger, with 56% stronger functional sensitivity (0.064 vs. 0.041) and 61% higher SNR. These results demonstrate that backscattered intensity-weighted mapping enhances sensitivity in both cortical and deep-brain regions while preserving super-resolution detail. Although both methods remain limited by localization constraints—where closely spaced MBs may not be fully resolved—the incorporation of backscattering amplitude enables B-fULM to effectively compensate for under-sampling. Together, these results establish B-fULM as a practical and sensitive approach for volumetric functional neuroimaging, with potential applications in probing neurovascular coupling, mapping functional connectivity, and investigating brain-wide hemodynamic responses at microscopic scales.

However, certain limitations remain. First, B-fULM improves sensitivity by incorporating MB amplitude. However, fluctuations in MB infusion concentration can bias the functional signal. In addition, the *in vivo* experiment was performed under urethane anesthesia, which may alter neurovascular coupling and impact the observed functional responses. Second, the proposed 4-system, 1024-channel Verasonics NXT platform enabled continuous 500 Hz volumetric imaging, but the hardware is costly and technically complex. Broader adoption will require streamlined acquisition schemes that achieve similar volume rates using single-system or clinical ultrasound systems. Finally, our sensitivity model assumes linear superposition of per-bubble backscatter (*I* ∝ *N*_true_). In practice, MB response and post-processing can deviate from linearity due to saturation, attenuation or destruction at higher acoustic pressure or concentration, and log-compression. These nonlinear effects may reduce the linearity between amplitude and true MB concentration. Future work will focus on incorporating advanced nonlinear models to better characterize amplitude-based sensitivity.

## V. CONCLUSION

In this work, we presented backscattering functional ultrasound localization microscopy (B-fULM) and the first implementation of 3D fULM enabled by a synchronized 4-NXT systems, 1024-channel acquisition platform. A new statistical framework and 3D microbubble advection simulations predicted the sensitivity advantages of backscattered amplitude integration over conventional count-based fULM, particularly under high concentrations. *In vivo* rat brain experiments confirmed stronger, more robust stimulus-evoked responses for B-fULM while preserving spatial resolution. Together, these results establish B-fULM as a practical and high-sensitivity framework for super-resolved 3D functional ultrasound imaging

